# Chemical inhibition of SUMOylation activates the FSHD locus

**DOI:** 10.1101/2025.08.01.668136

**Authors:** Alice Nordlinger, Loéva Morin, Alexandra Andrieux, Jean Philippe Trani, Pierre Perrin, Nathalie Eudes, Anne Dejean, Frédérique Magdinier

**Affiliations:** Nuclear Organization and Oncogenesis Unit, Department of Cell Biology and Infection, Institut Pasteur, Université Paris Cité, 75015 Paris, France; INSERM, U993, 75015 Paris, France; Aix Marseille Univ, INSERM, Marseille Medical Genetics, 13005 Marseille, France

**Keywords:** SUMOylation, SUMO, SMCHD1, FacioScapuloHumeral Dystrophy, D4Z4, DUX4, muscle

## Abstract

Facioscapulohumeral muscular dystrophy (FSHD) is a progressive and debilitating muscle disease for which no cure currently exists. In the majority of cases, FSHD is associated with the contraction of the D4Z4 macrosatellite repeat array at the 4q35 locus, leading to the inappropriate activation of *DUX4*, normally expressed during early embryogenesis. In FSHD, the genetic contraction is accompanied by hypomethylation of the D4Z4 array. Although a connection between DNA hypomethylation and *DUX4* expression has been suggested, the precise mechanisms that regulate *DUX4* transcription remain incompletely defined. The post-translational modification by SUMO was shown previously to repress the expression of *Dux*, the *DUX4* homolog, in mouse embryonic stem cells. Based on these findings, we explored here the contribution of SUMOylation in the regulation of *DUX4* in human muscle cells. We demonstrate that TAK⍰981 (subasumstat), a selective SUMOylation inhibitor, promotes transcriptional reprogramming of the 4q35 locus and induces *DUX4* expression. Importantly, this activation occurs independently of changes in DNA methylation or SMCHD1 ATPase activity. Our findings identify SUMOylation inhibition as a novel regulatory pathway driving *DUX4* expression. This work uncovers the importance of SUMOylation in the epigenetic control of the 4q35 locus and *DUX4* transcription, providing a potential therapeutic strategy to modulate *DUX4* expression in FSHD.

## Introduction

SUMOylation is a reversible post-translational modification that regulates protein functions through covalent attachment of small ubiquitin-like modifier (SUMO) proteins to their protein substrates in eukaryotic cells ^1,2^. SUMO proteins exist as three main paralogues in mammals, SUMO1, SUMO2 and SUMO3, with SUMO2 and 3 sharing high similarities. SUMOylation is a dynamic energy-dependent process involving unique E1 and E2 enzymes and a series of E3 ligases. The reversible aspect of SUMOylation is ensured by action of specific SUMO proteases called SENPs (Figure 1a). The SUMO pathway, which predominantly targets transcription factors and chromatin-associated proteins ^3-6^, is a key regulator of nuclear processes and cell identity. In particular, SUMOylation was reported to regulate the expression of *Dux* in mESCs, with loss of SUMOylation leading to derepression of *Dux* and transition to a totipotent-like state ^7-10^. In mESCs, SUMOylation-dependent silencing of *Dux* was shown to involve Prc1.6 and Kap/Setdb1 recruitment onto the *Dux* locus ^7^, as well as proper Dppa2/Dppa4 ^8,9^ and histone H1 SUMOylation^10^.

**Figure 1:**
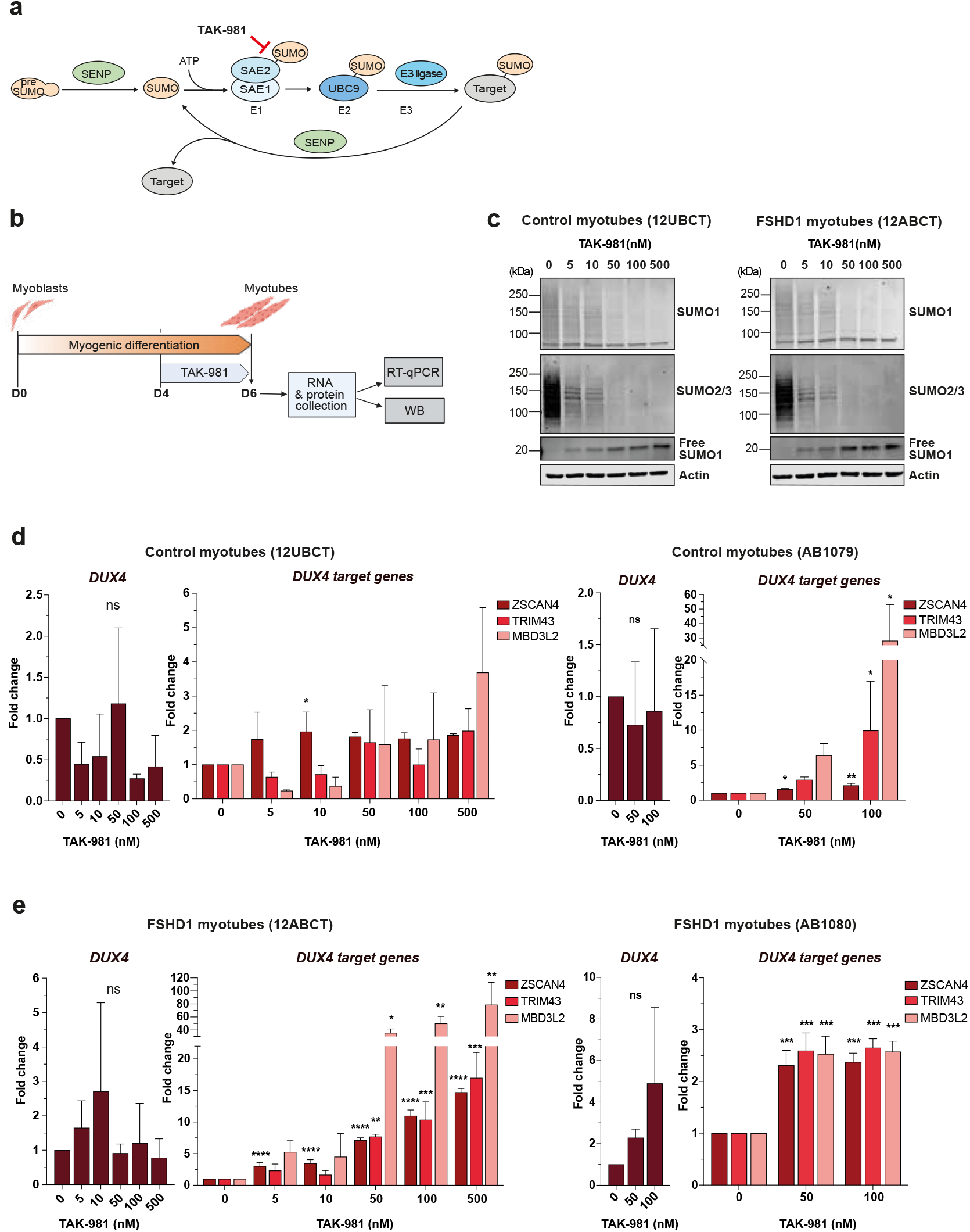
Chemical inhibition of SUMOylation induces a coordinated DUX4 program in FSHD1 patient-derived myotubes. **(a)** Representation of the reversible SUMOylation process. Before the first conjugation, a nascent SUMO precursor is proteolytically processed by a SUMO protease (SENP). Mature SUMO is then conjugated to a SAE1-SAE2 complex (E1) in an ATP-dependent manner. Next, activated SUMO is transferred to the E2 conjugating enzyme UBC9, before being conjugated to a target protein by the E3 ligase. SUMO conjugates can be cleaved from the target protein by a SENP, ensuring a dynamic and reversible SUMOylation process. TAK-981 inhibits SUMOylation by blocking activated SUMO at the catalytic site of SAE2. **(b)** Schematic representation of the experimental design for human hTERT-immortalized myoblasts. At confluence, control or FSHD1 hTERT-immortalized myoblasts were differentiated for 6 days in the presence of 2% horse serum and treated with DMSO or TAK-981 at different concentrations from day 4 to 6 of differentiation. At day 6, RNA and proteins were collected for gene expression analysis by RT-qPCR and western blot experiments. **(c)** Immunoblots for SUMO1 and SUMO2/3 in myotubes from control (12UBCT) and FSHD1 (12ABCT) cells. Actin was used as a loading control. **(d)** *DUX4* and *DUX4* target gene expression analyses in two cell lines of control myotubes (12UBCT and AB1079). Fold-changes were calculated relative to three housekeeping genes and normalized to the untreated (0) condition for each gene. Error bars represent mean ± SD. n=3. Significance: one-way ANOVA testing. ns: not significant. ^*^ p<0.05, ^**^ p<0.01, ^***^ p<0.001 and ^****^ p<0.0001. **(e)** *DUX4* and *DUX4* target gene expression analyses in myotubes obtained from two FSHD1 patients (12ABCT and AB1080). Fold-changes were calculated relative to three housekeeping genes and normalized to the untreated (0) condition for each gene. Error bars represent mean ± SD. n=3 independent experiments. Significance: one-way ANOVA testing. ns: not significant. ^*^ p<0.05, ^**^ p<0.01, ^***^ p<0.001 and ^****^ p<0.0001

The mouse Dux and human DUX4 transcription factors are involved in the zygotic genome activation (ZGA) during the 2C-4C stage in mouse and 4C-8C stage in human embryos, respectively ^11,12^. In human cells, DUX4 is encoded by the D4Z4 macrosatellite element that is located in the subtelomeric region of the 4q arm (4q35 locus) ^13^. Each D4Z4 element is approximately 3.3 kilobases in size and forms large tandemly repeated array structures at the subtelomeric 4q35 and 10q26 loci ^14,15^. Smaller size or partial D4Z4-like sequences are scattered in heterochromatin regions throughout the genome, in particular at the p arm of acrocentric chromosomes. At 4q35 and 10q26 loci, the number of repeat units can vary between individuals, typically ranging from 11 to over 100 ^16^. These arrays are highly GC-rich and present with heterochromatin features such as DNA methylation, leading to epigenetic repression ^17-19^.

The 4q35 locus is linked to type 1 Facioscapulohumeral Dystrophy (FSHD1, OMIM 158900), an autosomal dominant neuromuscular disease that is characterized by a progressive weakness and wasting of specific skeletal muscles of the face, shoulder girdle, and upper arms ^20,21^. In individuals with FSHD1, the D4Z4 repeat array on chromosome 4q35 is contracted to contain 1 to 10 repeat units, compared to the normal size range of 11-100 repeats in the unaffected population. This contraction is associated with D4Z4 DNA hypomethylation and chromatin opening enabling aberrant expression of the *DUX4* gene from the most distal D4Z4 repeat. Importantly, disease manifestation requires the presence of a permissive 4qA haplotype distal to the last D4Z4 unit, which provides a functional polyadenylation signal necessary for stable *DUX4* transcript production ^22^. FSHD2 (5% of patients, OMIM #158901) is clinically similar to FSHD1 but arises from a digenic mechanism in the absence of D4Z4 array contraction. In approximately 80% of these FSHD2 patients, the disease is associated with pathogenic variants in *SMCHD1* and involves D4Z4 hypomethylation ^23,24^, allowing *DUX4* expression from a normally sized repeat array ^25^. In human somatic cells, experimental overexpression of *DUX4* activates the expression of more than 400 genes. This cascade of transcriptional activation has been associated with immune response, muscle inflammation and atrophy or inhibition of muscle regeneration ^26^.

Given the role of SUMOylation in the regulation of *Dux* in mESCs, we decided to evaluate whether the SUMO pathway may similarly regulate the expression of *DUX4* in human muscle cells and, in particular, in the context of FSHD. To this aim, we exploited two different cellular models in which we decreased the global cellular levels of SUMOylation using a selective chemical inhibitor to investigate the role of SUMOylation in the regulation of *DUX4* and the 4q35 region, as well as the possible involvement of SMCHD1 post-translational modifications ^27^. We show that TAK⍰981 (subasumstat), a first-in-class SUMOylation inhibitor^28^, promotes transcriptional reprogramming at the 4q35 locus and induces *DUX4* expression, independently of changes in DNA methylation or SMCHD1 ATPase activity.

## Results

### SUMOylation inhibition activates the *DUX4* program in human myotubes

Given the repressive role of SUMOylation on *Dux* expression in mESCs, we explored the consequences of loss of SUMOylation in human muscle cells, in particular in the context of FSHD, which is characterized by aberrant expression of *DUX4* in skeletal muscle. To this end, we treated myotubes derived from hTERT-immortalized control and FSHD myoblasts with the TAK-981 SUMOylation inhibitor (henceforth called SUMOi) (Figure 1a) and analyzed the expression of *DUX4* and some of its target genes by RT-qPCR. TAK-981 ^28^ covalently binds to the SUMO-E1 Activating Enzyme complex (SAE1) complex and prevents the activation and transfer of SUMO proteins to SUMO E2 UBC9 and downstream protein targets (Figure 1a). hTERT-immortalized myotubes were treated for 48 hours (from day 4 to day 6 post-differentiation) with increasing doses of TAK-981 (0-500nM, final concentration) and harvested for molecular analyses (Figure 1b).

By western blotting, we observed a progressive and dose-dependent accumulation of free SUMO coupled with a decrease in SUMO1- and SUMO2/3-conjugated proteins in control and FSHD myotubes (Figure 1c, Figure S1a,b), attesting the efficacy of TAK-981 at tested concentrations. We next showed that SUMOylation inhibition is associated with a variegated induction of *DUX4* expression, mainly in FSHD myotubes (Figure 1d, e, Figure S1c). Concomitantly, we observed a slight dose-dependent increase in the expression of some of the selected DUX4 target genes (*ZSCAN4, TRIM43, MBD3L2*) in control cells (12UBCT ^29^, AB1079). Increase in DUX4 target gene expression is more pronounced and statistically significant in FSHD cells (12ABCT ^29^, AB10180), in particular for high doses of TAK-981 (Figure 1e). In addition, we noted a moderate but significant impact of SUMOi on the expression of the *MyoG* myogenic marker in both FSHD and control cells, with a maximal effect at higher doses of TAK-981 (Figure 2a, b). We also reported a dose-dependent decreased expression in *MyH2* and *MyH3* together with a decreased in the proportion of myotubes in FSHD cells compared to controls (Figure 2c).

**Figure 2:**
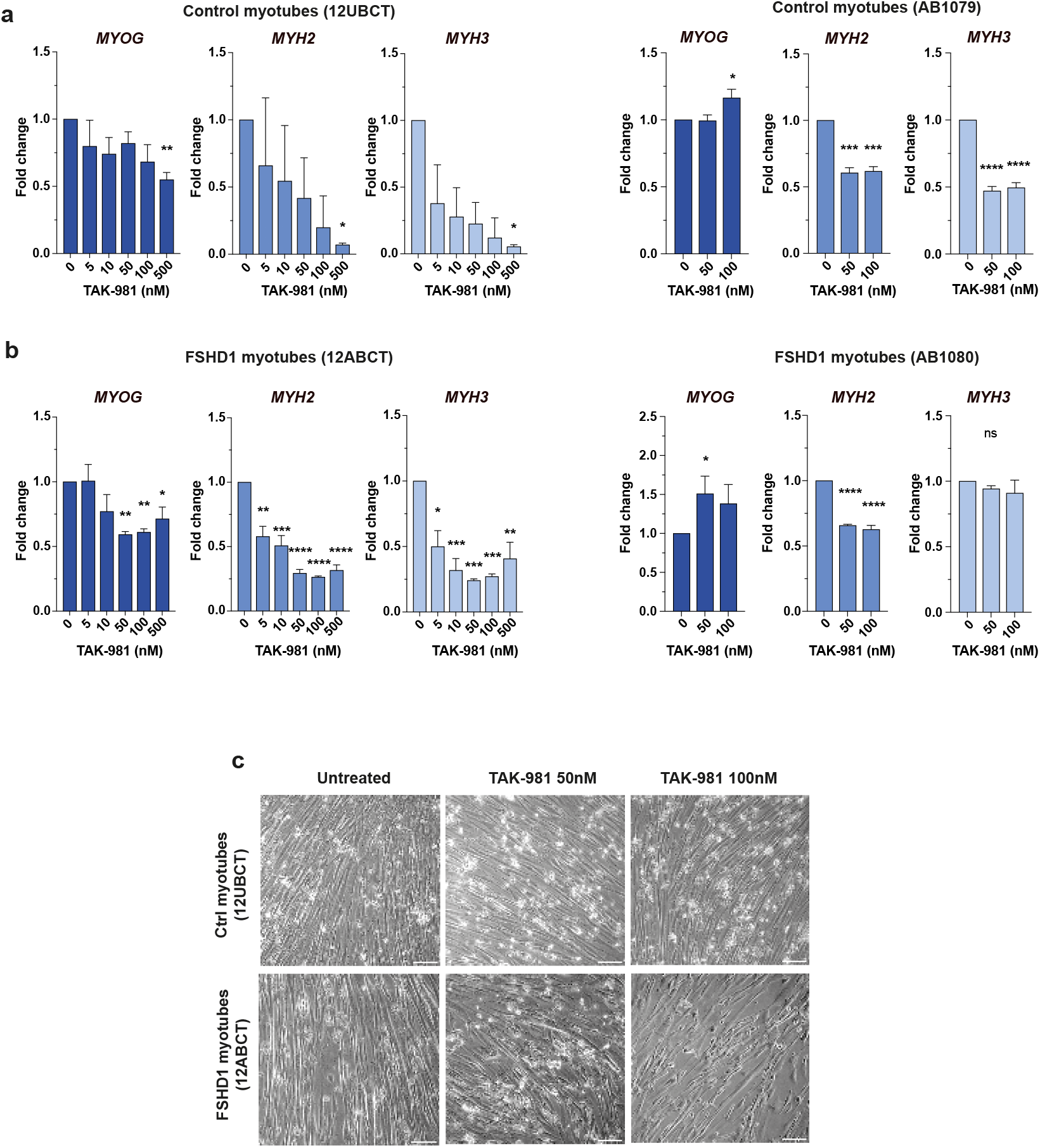
Inhibition of SUMOylation impairs myogenic gene expression in control and FSHD1-patient derived myotubes. **(a)** *MYOG, MYH2* and *MYH3* expression analyses in control myotubes (12UBCT and AB1079). Fold-changes were calculated relative to three housekeeping genes and normalized to the untreated (0) condition for each gene. Error bars represent mean ± SD. n=3. Significance: one-way ANOVA testing. ns: not significant. ^*^ p<0.05, ^**^ p<0.01, ^***^ p<0.001 and ^****^ p<0.0001 **(b)** *MYOG, MYH2* and *MYH3* expression analyses in myotubes obtained from FSHD1 patients (12ABCT and AB1080). Fold-changes were calculated relative to three housekeeping genes and normalized to the untreated (0) condition for each gene. Error bars represent mean ± SD. n=3. Significance: one-way ANOVA testing. ns: not significant. ^*^ p<0.05, ^**^ p<0.01, ^***^ p<0.001 and ^****^ p<0.0001 **(c)** Brightfield images of differentiated control and FSHD1 myotubes at day 6 after TAK-981 treatment at indicated concentrations (day 4-6). Scale bar = 50µm.

### TAK-981 enhances *DUX4* expression via SUMOylation inhibition in hiPSC-derived muscle cells

Next, we investigated the impact of SUMOi on *DUX4* expression during muscle tissue differentiation. To this aim, we treated control and FSHD muscle cells derived from induced pluripotent cells (hiPSCs) over a period of 15 days post-differentiation with daily change in cell culture medium supplemented with TAK-981 or DMSO alone (Figure 3a). Different concentrations of TAK-981 were used and cells were collected at the end of the differentiation procedure (day 30, D30) for RNA and protein extraction. Efficiency of SUMOylation inhibition was verified by western blotting (Supplementary Figure 2a-c).

**Figure 3:**
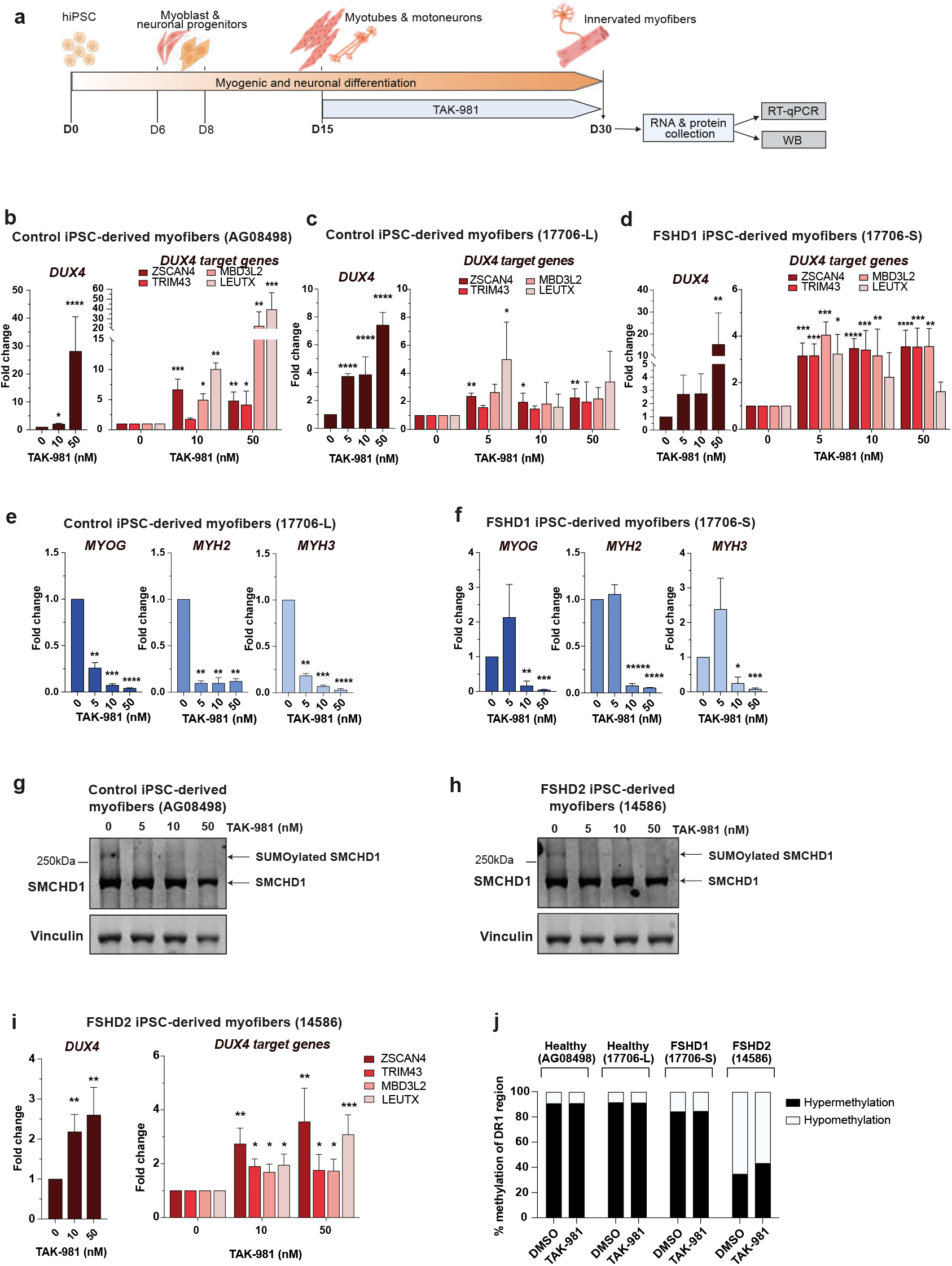
Loss of SUMOylation triggers DUX4 activation in FSHD1 iPSC-derived myotubes. (**a**) Experimental design for hiPSC-derived myofibers. HiPSCs derived from a healthy donor or FSHD1 patient were differentiated for 30 days as described^50,51^. TAK-981 was added every day from day 15 to 30. Cell culture medium supplemented in DMSO or TAK-981 was changed every 24 hours. At day 30, RNA and proteins were collected. (**b**-**d**) *DUX4* and *DUX4* target gene expression analyses in control AG08498 (**b**), control 17706-L (**c**) or FSHD1 17706-S (**d**) hiPSC-derived myofibers. Healthy iPSC (17706-L) and FSHD1 iPSC (17706-S) are isogenic clones obtained from a mosaic patient (17706). Fold-changes were calculated relative to three housekeeping genes and normalized to the untreated (0) condition for each gene. Error bars represent mean ± SD. n=3. Significance: one-way ANOVA testing. ns: not significant. ^*^ p<0.05, ^**^ p<0.01, ^***^ p<0.001 and ^****^ p<0.0001 (**e**-**f**) *MYOG, MYH2* and *MYH3* expression analyses in the control 17706-L (**e**) and FSHD1 17706-S (**f**) hiPSC-derived myofibers. Fold-changes were calculated as above. Error bars represent mean ± SD. n=3. Significance: one-way ANOVA testing. ns: not significant. ^*^ p<0.05, ^**^ p<0.01, ^***^ p<0.001 and ^****^ p<0.0001 (**g**-**h**) Immunoblots for the endogenous SMCHD1 protein in control AG08498 (**g**) and FSHD2 14586 **(h)** hiPSC-derived myofibers treated with TAK-981 at indicated concentrations. Vinculin is used as a loading control. Arrows indicate SMCHD1 and its SUMOylated form. **(i)** *DUX4* and *DUX4* target gene expression analyses in the FSHD2 hiPSC-derived myofibers (14586). Fold-changes were calculated as above. Error bars represent mean ± SD. n=3. Significance: one-way ANOVA testing. ns: not significant. ^*^ p<0.05, ^**^ p<0.01, ^***^ p<0.001 and ^****^ p<0.0001 **(j)** Percentage of methylated (black) and unmethylated (white) CpG at the DR1 region located in the proximal part of the D4Z4 repeat (*y-axis*). Methylation was determined for each CpG within the sequence of interest. Each bar represents the percentage of methylated or unmethylated CpG for each CpG (*x-axis*). The global methylation level was determined for all sequenced clones and all CpG in the region of interest after sodium bisulfite sequencing analysis in control and FSHD hiPSC-derived myofibers after TAK-981 or DMSO treatment.

In control cells, we observed a robust increase in *DUX4* expression at the highest dose of TAK-981 (Figure 3b) associated with increased expression of selected DUX4 target genes, *MBD3L2* and *LEUTX*, in particular. For FSHD1, we first compared the effect of SUMOi on the transcriptional activation of *DUX4* and DUX4 target gene in hiPSC-derived muscle tissue for a mosaic patient for which healthy (Figure 3c) and diseased (Figure 3d) isogenic clones were isolated ^30^. In both contexts, we observed a dose-dependent increase in *DUX4* expression together with a significant increase in the expression of DUX4 target genes that is more consistent in FSHD cells (17706-S) and observed at low doses of TAK-981 (Figure 3c, d). As showed for immortalized myoblasts, the expression of the *MyoG* myogenic factor and of Myosin genes (*MyH2, MyH3*) was decreased in both control and FSHD cells, at low doses of TAK-981 (Figure 3e,f).

### Impact of loss of SUMOylation on SMCHD1 and SMCHD1-deficient cells

The SMCHD1 protein contains six principal lysine residues (K784, K1373, K1496, K1958, K1976, K2002) modified by SUMO2/3^27^. In hiPSC-derived myofibers, SUMOylated SMCHD1 was detected as an upper band by western blotting (Figure 3g,h). This upper band is no longer detected after TAK-981 treatment in both controls and FSHD cells, showing that SMCHD1 SUMOylation is decrease upon TAK981 treatment.

We next assessed whether SMCHD1 loss of function in SMCHD1-deficient cells together with loss of SUMOylation might exacerbate *DUX4* induction. We treated FSHD2 hiPSC-derived muscle cells from D15 to D30 of differentiation with SUMOi using hiPSC from a FSHD2 patient carrying a pathogenic *SMCHD1* that abrogates the protein ATPase activity ^30^ (Figure 3i). *DUX4* and its target genes exhibited levels of expression similar to those found in FSHD1 patient-derived myofibers (Figure 3d,i). In addition, the presence of a loss of function variant in *SMCHD1* does not seem to affect SUMOylation of the protein in basal conditions in FSHD2 hiPSC-derived myofibers (Figure 3h). We concluded that SUMOylation depletion promotes *DUX4* expression in both control and FSHD cells, regardless of SMCHD1 catalytic activity.

### SUMOylation inhibition does not impact D4Z4 methylation profile

As the D4Z4 macrosatellite is differentially methylated at the DNA level between non-affected individuals and FSHD patients, we next investigated whether *DUX4* induction upon TAK-981 resulted from D4Z4 hypomethylation. To this aim, hiPSC-derived muscle of two controls (healthy donor and healthy clone isolated from the mosaic patient), one FSHD1 patient (diseased clone from the mosaic patient) and one FSHD2 patient were exposed to the SUMOylation inhibitor from D15 to D30 post-differentiation as described (Figure 3a). Genomic DNA was extracted at the end of the treatment and D4Z4 methylation level was analyzed by sodium bisulfite sequencing at the DR1 region which is hypomethylated in FSHD patients compared to non-affected individuals ^23,24^. In all conditions, we compared cells treated at a final concentration of 50nM TAK-981 to cells treated with DMSO only. Inhibition of SUMOylation does not impact the level of methylation at DR1 in all tested hiPSC derived-myofibers (Figure 3j), indicating that D4Z4 chromatin relaxation leading to TAK-981-induced activation of *DUX4* expression does not involve the hypomethylation of the repeat.

### Loss of SUMOylation modulates expression of genes at the 4q35 locus

We then evaluated whether TAK-981-induced hypoSUMOylation only affects *DUX4* expression or also the expression of other genes at the 4q35 locus. We analyzed expression of four different genes within a 7Mb range of the D4Z4 region (*WWC2*) and located in different Topologically Associated Domains (TADs) ^31^. We included genes previously implicated in FSHD such as *FRG2* ^32-34^, *FRG1* ^32^ and *FAT1* ^35^ (Figure 4a). The expression of the most distal gene *WWC2* is not impacted by TAK-981 treatment compared to mock-treated controls or FSHD immortalized myotubes (Figure 4b,c). Conversely, we observed a robust and statistically significant upregulation of *FRG2* and *FAT1* at low doses of TAK-981 in both control and FSHD1 cells, but a more heterogenous effect on *FRG1* expression depending on samples (Figure 4b-c).

**Figure 4:**
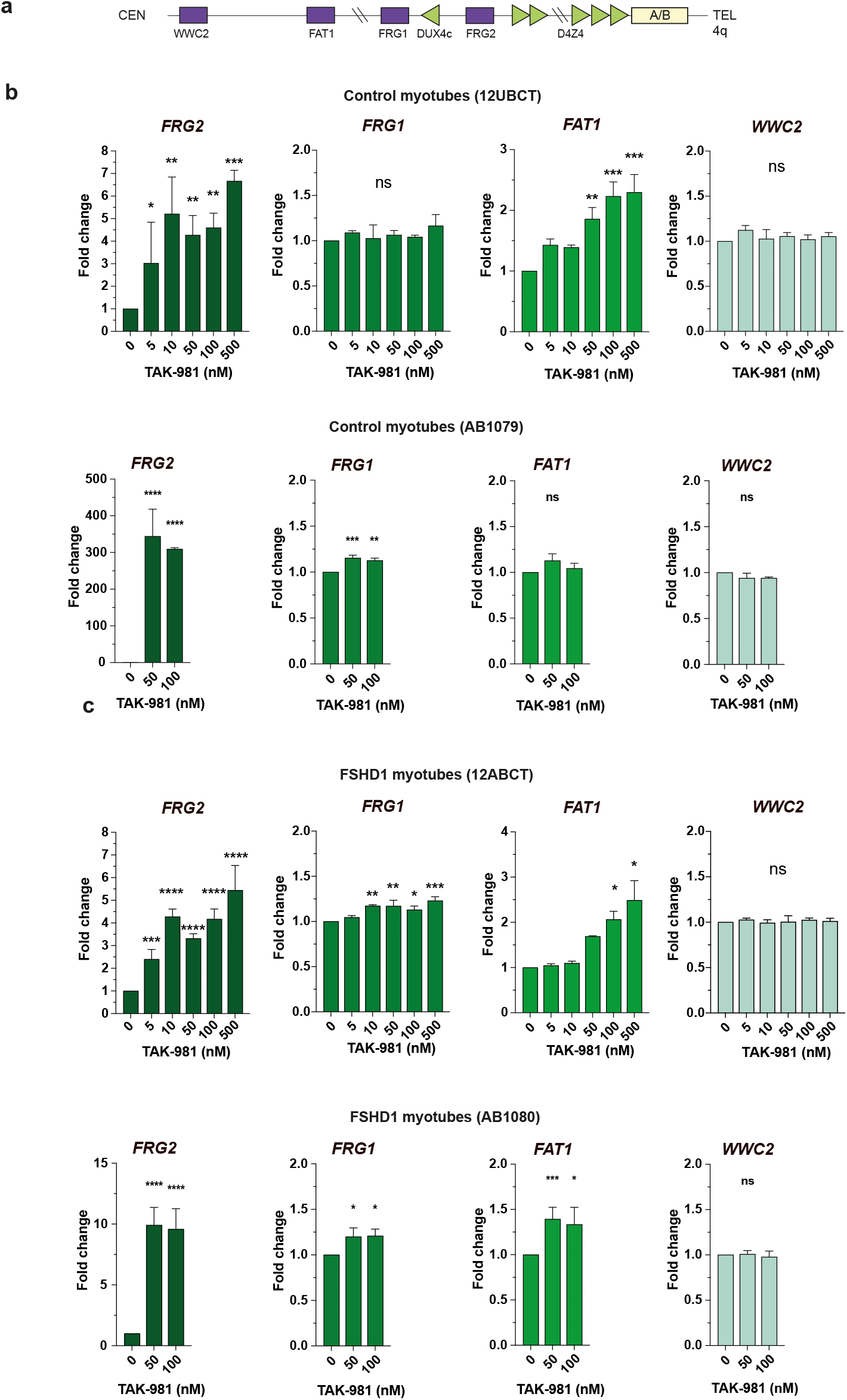
SUMOylation inhibition affects expression of other genes of the *DUX4* locus in myotubes. (**a**) Schematic representation of human chromosome region 4q35. (**b**-**c**) *FRG2, FRG1, FAT1* and *WWC2* expression analyses in control myotubes 12UBCT and AB1079 (**b**) and in FSHD1 patient-derived myotubes 12ABCT and AB1080 (**c**). Fold-changes were calculated relative to three housekeeping genes and normalized to the untreated (0) condition for each gene. Error bars represent mean ± SD. n=3. Significance: one-way ANOVA testing. ns: not significant. ^*^ p<0.05, ^**^ p<0.01, ^***^ p<0.001 and ^****^ p<0.0001

The same trend was visible in hiPSC-derived muscle tissue when cells were treated during differentiation, in particular for *FRG2* and *FAT1*, both in controls and FSHD cells (Figure 5a-d), suggesting that SUMOylation inhibition contributes to the relaxation of the 4q35 locus.

**Figure 5:**
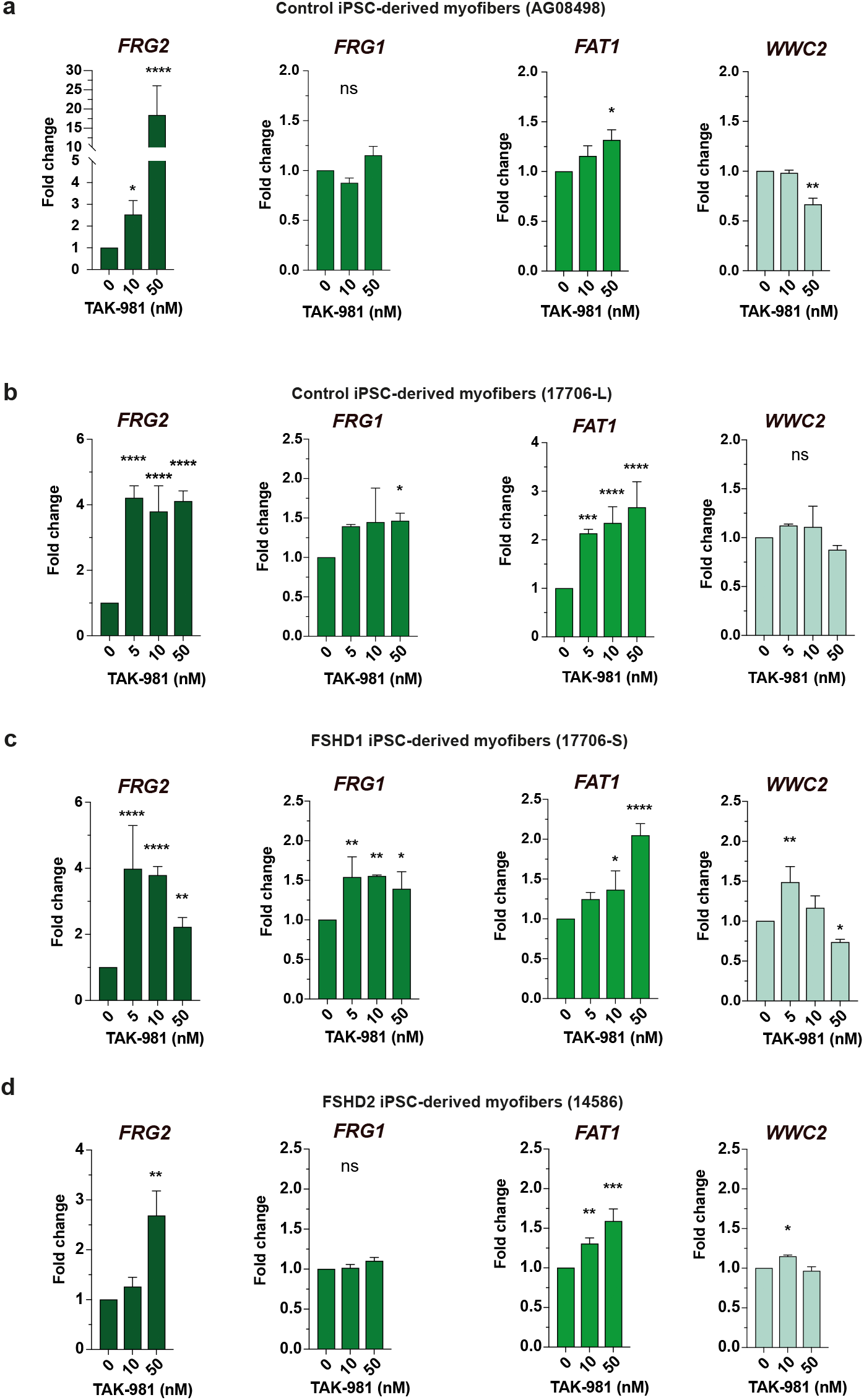
Expressions of genes in *DUX4* locus are affected in hiPSC-derived myofibers. (**a**-**d**) *FRG2, FRG1, FAT1* and *WWC2* expression analyses in control AG08498 (**a**), control 17706-L (**b**), FSHD1 17706-S (**c**) and FSHD2 14586 (**d**) hiPSC-derived myofibers. Fold-changes were calculated relative to three housekeeping genes and normalized to the untreated (0) condition for each gene. Error bars represent mean ± SD. n=3. Significance: one-way ANOVA testing. ns: not significant. ^*^ p<0.05, ^**^ p<0.01, ^***^ p<0.001 and ^****^ p<0.0001

## Discussion

Post-translational modification by SUMO acts to stabilize cell identity in a variety of contexts ^7,36,37 38^. In particular, hypoSUMOylation enhances pluripotency reprogramming *in vitro and in vivo* and suppression of SUMOylation promotes the spontaneous conversion of ESCs to 2-cell-embryo-like (2C-like) state^7,8,38^. This transition involves two distinct mechanisms. First, hypoSUMOylation activates transcription of genomic loci regulated by the H3K9 histone methyltransferases Setdb1 and Suv39H1 by inducing a global decrease in H3K9 trimethylation ^7,39-41^. Second, hypoSUMOylation leads to *Dux* derepression by interfering with the recruitment of Prc1.6 and Kap/Setdb1 at the *Dux* promoter^7^, together with impairing Dppa2/Dppa4 and histone H1 repressing activities^1,9,10^.

Mouse *Dux* and human *DUX4* both encode a transcription factor that plays a key role in triggering zygotic genome activation (ZGA) in the early stages of embryogenesis ^11,12^. In human somatic cells, *DUX4* is normally silenced through epigenetic repression of the D4Z4 macrosatellite repeat array on chromosome 4q35, involving DNA methylation and repressive histone marks such as H3K9me3 and H3K27me3 ^22,42^. In FSHD, ectopic expression of *DUX4* has been proposed as the main trigger leading to muscle dysfunction ^22^. Activation of *DUX4* in a subset of muscle cells is subsequent to the reduction in the number of D4Z4 macrosatellite at the 4q35 locus in FSHD1 ^22^ or to the presence of a variant in *SMCHD1* in FSHD2 ^25^, both leading to hypomethylation of D4Z4 and chromatin relaxation of the 4q35 locus. However, the mechanisms underlying burst of *DUX4* transcriptional activation in muscle remain unknown.

A large number of transcription factors and chromatin regulators are SUMOylated, and SUMOylation is generally associated with transcriptional repression ^3-6,27^. SUMO facilitates the recruitment of SUMO interacting motif (SIM)-containing proteins ^43,44^ such as histone deacetylases or the Setdb1 histone methyltransferase. Interestingly, the SMCHD1 protein contains six main lysine residues (K784, K1373, K1496, K1958, K1976, K2002) that are modified by SUMOylation and 15 minor acceptor lysines containing the consensus sequence required for SUMOylation ^27^. Several of these arguments point towards a role for SUMOylation in the regulation of the 4q35 locus and potentially of aberrant *DUX4* expression in the context of FSHD. However, nothing is currently known on how SUMOylation might regulate SMCHD1 function.

Recent development of a highly specific small molecule inhibitor of SUMOylation, TAK-981, proved to be a powerful tool for studying various facets of SUMOylation biology ^28^. Notably, TAK-981 is able to induce a strong spontaneous type I interferon response in myeloid cells^45^, mirroring genetic inactivation of SUMOylation^46^. In line with this finding, TAK-981 was shown to activate antitumor immune response in pre-clinical models^45,47^ and in the clinics ^48^. Here, we show that short or prolonged treatment of muscle cells *in vitro* with TAK-981 activates the *DUX4* expression program in control cells and exacerbates *DUX4* activation in cells from FSHD patients. Using cells from a patient with FSHD2 carrying a variant in *SMCHD1* altering the functional activity of the ATPase domain and showing a marked decrease in D4Z4 methylation ^30^, we showed that the combination of SMCHD1 loss of function and D4Z4 hypomethylation does not lead to increased TAK-981-induced *DUX4* activation beyond levels observed in TAK-981-treated FSHD1 cells. We concluded that D4Z4 hypomethylation alone is not sufficient to control *DUX4* expression.

In addition, we showed that treatment with SUMOi increases expression of other genes located at the 4q35 locus such as the *FRG2* non-coding RNA and *FAT1*, that might also contribute to the FSHD phenotype ^32-35^. This suggests that SUMOylation inhibition contributes to a more global relaxation of genes located in the vicinity of the D4Z4 macrosatellite array, in particular genes located within the same TAD and showing variable long-distance interaction depending on the cell context ^31^.

Notably, recent research by an independent group investigated the link between SUMOylation and *DUX4* regulation in FSHD, using ML-792, an alternative inhibitor of the SUMO-conjugation pathway in cultured cells ^49^. As observed in our cell models, the authors demonstrated that SMCHD1 is SUMOylated in both myoblasts and myotubes. Consistent with our observations, treatment with ML-792 induces *DUX4* expression and a variable decrease in myogenic gene expression. Their analysis of SMCHD1 mutants lacking SUMOylation sites revealed no significant impact on SMCHD1 protein stability, nuclear localization, dimerization, or chromatin binding at the D4Z4 locus. Similarly, our results on FSHD2-patient derived cells with a loss-of-function of SMCHD1 did not uncover a definitive role for SUMOylated SMCHD1 in regulating the 4q35 region, indicating that further functional studies are needed ^49^.

Overall, we provide here a proof of concept of the effect of SUMOylation in the regulation of *DUX4* expression and epigenetic regulation of the 4q35 region induced by a second-generation SUMO E1 inhibitor, TAK-981, in different models of control and FSHD muscle cells. Our data suggest that manipulating SUMOylation, potentially through SENP deSUMOylase inhibition, may help silencing inappropriate de-repression of DUX4, opening new avenues for the development of therapeutic strategies targeting SUMOylation in FSHD. Further studies will be essential to define the therapeutic relevance of such approaches.

## Acknowledgments

We are grateful to Agnès Marchio for help in the RT-qPCR settings and Jacob Seeler for helpful discussions.

