## Supplemental information for "Chemical inhibition of SUMOylation activates the FSHD locus"

**Supplementary information**

**Legends of Supplementary Figures**

**Supplementary Figure 1: Validation of TAK-981 induced hypoSUMOylation in hTERT immortalized myotubes**

(**a**) Immunoblots for SUMO1 and SUMO2/3 in control (AB1079) myotubes. Actin is used as a loading control.

(**b**) Immunoblots for SUMO1 and SUMO2/3 in FSHD1-patient derived myotubes (AB1080). Actin is used as a loading control.

(**c**) Electrophoresis of *DUX4* qPCR products in all tested control (12UBCT, AB1079) and FSHD1-patient derived myotubes (12ABCT, AB1080) treated with indicated concentrations of TAK-981 from day 4 to day 6 of differentiation.

**Supplementary Figure 2: Validation of TAK-981 induced hypoSUMOylation in hiPSC-derived muscle cells.**

(**a**-**d**) Immunoblots for SUMO1 and SUMO2/3 in control AG08498 (a), control 17706-L (b), FSHD1 17796-S (c) and FSHD2 14586 (d) hiPSC-derived myofibers. Actin is used as a loading control.

**Supplementary Table S1. List of primers used for RT-qPCR analysis**

| **Gene** | **Sequence Forward** | **Sequence Reverse** |
| --- | --- | --- |
| *HPRT* | TGATAGATCCATTCCTATGACTGTA | CAAGACATTCTTTCCAGTTAAAGTTG |
| *PPIA* | ATGCTGGACCCAACACAAAT | TCTTTCACTTTGCCAAACACC |
| *GUSB* | CCGAGTGAAGATCCCCTTTTTA | CTCATTTGGAATTTTGCCGATT |
| *DUX4* | AGGCGCAACCTCTCCTAGAAA | GCTCCTCCAGCAGAGCCCGGTATTC |
| *ZSCAN4* | TGGAAATCAAGTGGCAAAAA | CTGCATGTGGACGTGGAC |
| *TRIM43* | ACCCATCACTGGACTGGTGT | CACATCCTCAAAGAGCCTGA |
| *MBD3L2* | GCGTTCACCTCTTTTCCAAG | GCCATGTGGATTTCTCGTTT |
| *LEUTX* | CTTCAAAGCTACAACTTGATCTATCC | AGTCTCCTCCTTCTTCACTGA |
| *FRG2* | CGCACCTTTCACTTGAGCTT | GAATGGGAGAAGGCGGTCT |
| *FRG1* | TTGTTGGAATCTGGTGGACA | CCATTGTCGAGTGCATGTATATAGG |
| *FAT1* | CATTAGAGATGGCTCTGGCG | ATGGGAGGTCGATTCACG |
| *WWC2* | TGACAATATGGCAGTTCGCCCCA | TCACTGTCACTCCGATTTAACCTGC |
| *MYOG* | CAGCGAATGCAGCTCTCA | GGTTGTGGGCATCTGTAGG |
| *MYH2* | TTCTCAGGCTTCAAGATTTGG | CTGGAGCTTGCGGAATTTAG |
| *MYH3* | ATCGTGAAAACCAGTCCATTCT | TTGGCCAGGTCCCCAGTAGCT |

­

Material & Methods

**Cell lines and culture**

hTERT immortalized myoblasts (12UBCT, 12ABCT) were described in ^1^. The 12ABCT cells were obtained from a FSHD1 patient carrying 5 D4Z4 units). The hTERT-immortalized AB1080, AB1079 line were provided by the MyoLine platform of the Institute of Myology (Paris, France). The AB1079 cells line was derived from paravertebral muscles of a FSHD1 patient carrying 5 D4Z4 units. These cell lines were cultured in proliferative conditions in a Growth Medium composed of 4:1 volumes of Dulbecco's modified Eagle medium (DMEM, Gibco)/Medium 199 (Gibco), supplemented with 15% fetal bovine serum (Euroclone), 0.02 m HEPES buffer (Invitrogen), 1.4 mg L^−1^ vitamin B12 (Sigma); 0.03 mg L^−1^ ZnSO4 (Fisher Scientific), 0.055 mg L^−1^ dexamethasone (Sigma), 2.5 µg L^−1^ hepatocyte growth factor (Chemicon International) and 10 µg L^−1^ basic fibroblast growth factor (BioPioneer) in the presence of 1% Penicillin/streptomycin.

All hiPSC clones were generated from reprogramming of primary fibroblasts as described in^2^. The C5 hiPSC clone (AG08498) is derived from human fibroblasts purchased from the Coriell Institute^3^. The 17706 hiPSC clones, 17706-L (long) and 17706-S (short, 2 D4Z4 units) were derived from a mosaic patient carrying one normal 4q35 allele and one shorter allele with 2 RU (FSHD1; female; age 56 at sampling). The 14586 hiPSC clone was derived from a patient carrying two normal 4q35 alleles but one *SMCHD1* mutation (c.573A>C; p.Q193P; FSHD2; male; age 67 at sampling). For clones generated from patients, individual consents were provided for using their sample in medical research. hiPSC colonies were grown and expanded in mTeSR1 medium (Stemcell Technologies) on BD Matrigel (BD Biosciences, 354277) coated dishes as described^2,3^.

**Cell differentiation and pharmacological treatment**

For human immortalized myoblasts, once 90% confluency was reached, Growth Medium was replaced for 6 days by Differentiation Media (DM) composed of 4:1 DMEM/Medium 199 supplemented with 2% Horse Serum (Gibco) to induce final differentiation. Cells were treated with DMSO or TAK-981 (HY-111789, MedChemExpress) diluted in Differentiation Medium from day 4 to day 6 and changed every day.

Differentiation of hiPSC clones was performed as described ^4,5^. Briefly, at day 0 they were switched for a differentiation medium (neurobasal medium supplemented with N2 1X, B27 1X, penicillin/streptomycin 1X, non-essential amino-acid 1X, Glutamax 1X ; Life Technologies) changed every day, containing ITS-A 1X (51300044, Life Technologies), LDN193189 0,5µM (SML0559, Sigma-Merck) and CHIR99021 3µM (5ML-1046, Sigma-Merck). From day 6 to day 8, medium is replaced by a differentiation medium containing LDN193189, IGF1 4ng/mL, HGF 10ng/mL (100-39, Peprotech) and β-mercaptoethanol 90,8µM (31350010, ThermoFisher). At day 8, medium is changed for a differentiation medium supplemented only with IGF1and β-mercaptoethanol for four days. At day 12, a differentiation medium was used supplemented with IGF1, β-mercaptoethanol and DAPT 10µM (Tocris Biosciences) until day 17. From day 17 to the end of experiment at day 30, medium was changed every day for differentiation medium supplemented with IGF1. Cells were treated from day 15 to day 30 with DMSO or TAK-981 diluted in the appropriate differentiation medium and changed every day.

**Quantitative real-time PCR**

Total RNA was collected at the end of differentiation process at day 6 for human immortalized myoblasts and day 30 for hiPSC clones. Total RNA was extracted using TriZol reagent (Invitrogen) following manufacturer’s instructions and quantiﬁed using a spectrophotometer (ND-1000, Nanodrop Technologies). 2µg of RNA were used to synthetize cDNA using SuperScript IV kit (18091050, Invitrogen) preceded by a step of genomic DNase digestion with ezDNase (11766051, Invitrogen). Quantitative PCR was performed on a CFX96 real Time System Thermal Cycler (Bio-Rad) using SYBR Green PCR Mastermix (Applied Biosystems) and specific primers available in Table S1.

**Immunoblots**

Protein extracts were collected at the end of differentiation process at day 6 for human immortalized myoblasts and day 30 for hiPSC clones immediately in Laemmli buffer (Bio-Rad) 1X with β-mercaptoethanol to avoid rapid deSUMOylation of proteins. After boiling and sonication steps, protein dosage was performed using Ionic Detergent Compatibility Reagent for Pierce 660nm Assay (22660, Pierce) and a SkanIt (ThermoFisher) spectrophotometer at 660nm. Equal quantities of protein extracts were loaded and were verified after transfer (Bio-Rad) by a Ponceau staining on nitrocellulose membranes. For SUMO immunoblots, Bis-Tris polyacrylamide 4-12% gels (Bio-Rad) were used. For endogenous SMCHD1 immunoblots, homemade gels at 6.5% Bis-acrylamide were prepared.

Anti-SUMO1 (ab32058, Abcam; 1/1000), anti-SUMO2/3 8A2 (ab81371 Abcam; 1/1000), anti-Vinculin (ab18058; Abcam; 1/1000), anti-SMCHD1 (ab175235; Abcam; 1/666) were applied as primary antibodies, followed by the secondary antibody at a 1:10,000 dilution, IRDye 800CW goat anti-rabbit (926-32211, LI-COR) and IRDye 680LT goat-anti-mouse (926-68020, LI-COR). Immunoreactivity was detected with an OdysseyTM Infrared-Imaging System (LI-COR), according to the OdysseyTM Western Blotting Protocol.

**Brightfield microscopy images**

Images of human immortalized myoblast were made at day 6 of differentiation using a Zeiss fast observer microscope at a 10X magnification.

***DUX4* electrophoresis**

qPCR products resulted from *DUX4* RT-qPCR amplification were loaded on 2% agarose gels (G800802; Invitrogen) and imaged using Gel Doc XR System (Bio-Rad) device.

**Sodium bisulfite sequencing**

For bisulfite modification, 1 μg of genomic DNA was denatured for 30 minutes at 37°C in NaOH 0.4N and incubated overnight in a solution of 3M Sodium bisulfite pH5 and 10mM Hydroquinone using previously described protocol ^6^. Converted DNA was then purified using the Wizard DNA CleanUp kit (Promega) following manufacturer’s recommendation and precipitated by ethanol precipitation for 5 hours at -20°C. After centrifugation, DNA pellet was resuspended in 40μL of water and stored at -20°C until use. Converted DNA was amplified using the DR1 primers avoiding the presence of CpGs in the primer sequence in order to amplify both methylated and unmethylated DNA with the same efficiency^7^. After sequencing of the PCR fragments, the methylation level was calculated as described^8^ and corresponds to the global level of methylation for each biological sample in a given region calculated as the ratio of methylated CpG with the number of aligned CpG for all sequences and CpG positions for a given biological sample.

**Quantification and statistical analysis**

For RT-qPCR analysis, fold-change values of gene expression were obtained using the 2^-ΔΔCt^ method. Briefly, ΔCt values were first calculated relative to the average Ct of three housekeeping genes (*PPIA, HPRT, GUSB*), then normalized to the untreated condition (0).

Statistical analyses of RT-qPCR data were performed using ordinary one-way ANOVA on the relative ΔCt values with GraphPad/Prism 10, with validation of the bioinformatics and biostatistics hub of Institut Pasteur. When significancy is observed, multiple comparisons were conducted with a Dunnett T3 test.
