## Supplementary figures and images for "Chemical inhibition of SUMOylation activates the FSHD locus"

### Supplementary figure 1

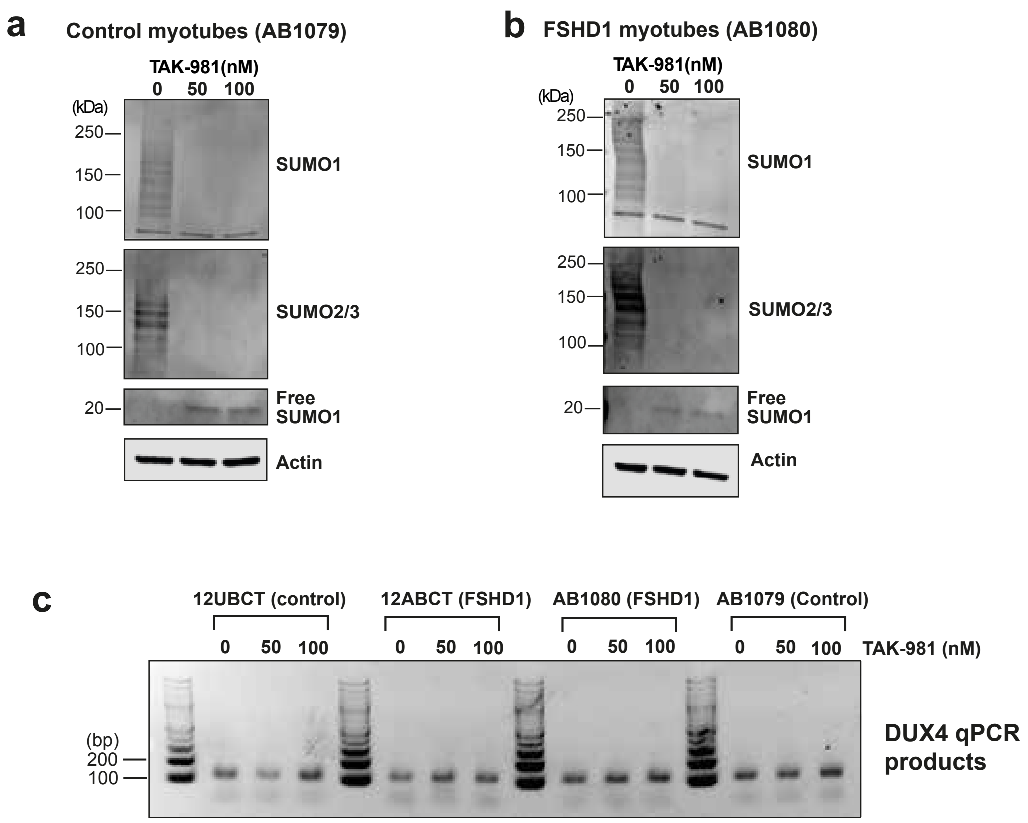


#### Supplementary figure 1

### Supplementary figure 2

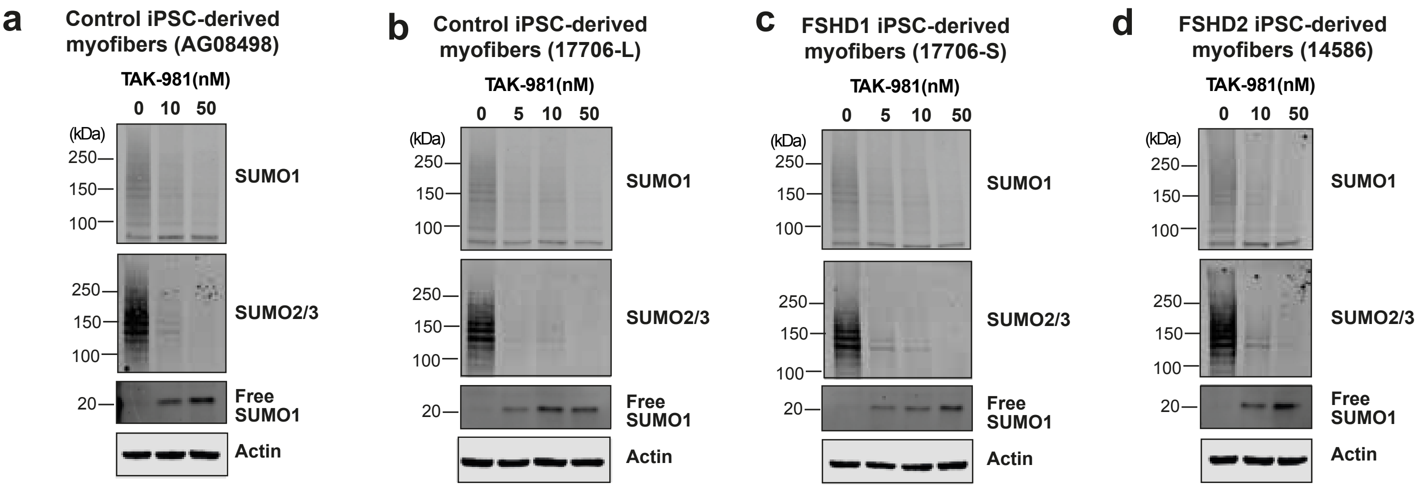


##### Supplementary Figure 2
